# Context Dependent Perturbation of Allelic Expression Imbalance Reveals Novel Candidate Therapeutic Targets for Metabolic diseases

**DOI:** 10.1101/2023.11.06.565672

**Authors:** Sunjin Moon, So-Young Park

## Abstract

**Background:** Obesity is a pivotal trigger for a spectrum of complex metabolic disorders. By colocalizing cis-eQTLs in adipose tissues from the GTEx consortium and trait-associated SNPs for complex traits from the GWAS Catalog within 3.6 million DNase I hypersensitive sites (DHSs), we systematically identify regulatory variants and genes that exhibit cis effects, as well as potential causal variants within the context of regulatory elements.

**Results:** Our analysis reveals that 229,504 (26.4%) cis-eQTLs located within DHS reside densely near the transcription start sites, contrasting with those outside of DHS. We observed that genes with higher allelic imbalance have shorter transcript lengths with larger number cis-eQTLs within DHS, and such imbalance genes are predominantly linked to signaling and immune response, whereas those with lower allelic imbalance tend to be involved in metabolism. Our composite colocalization score prioritizes 5,202 DHSs that encompass both cis-eQTLs and trait-associated SNPs, targeting 2,232 protein-coding genes and 523 lncRNAs across complex traits. We highlight the lncRNA SNHG5 as a prime example; it is associated with high-density lipoprotein levels and exhibits low allelic imbalance, and is also down-regulated in adipose tissue from patients with obesity.

**Conclusions:** Our findings underscore the critical role of regulatory context in pinpointing causal variants and refining target genes, offering rich insights into the genetic mechanisms pertinent to obesity and providing valuable resources for the diagnosis and therapeutic targeting of metabolic diseases.

## INTRODUCTION

Recent advances in the integration of various data sources enable better gene prioritization[1]. The advances in technology, such as deep sequencing, paired with the accumulation of large-scale omics data and complemented by refined analytical tools like advanced statistical methods, continue to expand our understanding in cell type- and disease-specific regulatory variant mapping [2–6]. Although colocalization offers promising insights, discerning causality from coincidence remains a challenge due to intricate linkage disequilibrium (LD) patterns. Discernment is crucial, reminding us that colocalization doesn’t automatically indicate causality and one should be cautious when interpreting these findings [7]. This challenge is further magnified by the effects of LD in methods such as identifying regulatory variants associated with complex traits and diseases in transcriptome-wide association studies [8,9].

Allelic-specific expression (ASE) remains one of the most direct ways to identify regulatory variants, where cis-acting regulatory elements play a fundamental role in chromatin structural alterations and DNA methylation [10,11]. Regulatory variants modulate allele-specific TF binding, DNase I hypersensitivity, and histone modifications, further shaping allelic expression imbalance [12]. Distinct expression heterogeneity between individuals is observed among genes affecting various biological functions[13–16]. Allelic imbalance (AI), a phenomenon where one allele of a heterozygous gene pair is preferentially expressed over the other, provides a nuanced yet essential layer of gene regulation that can contribute to individual variation in disease susceptibility.

The concept of allelic imbalance has evolved considerably over the last few years, AI lies at the epicenter of genetic phenomena, conserved strikingly across organisms[17–19]. From an evolutionary perspective, AI showcases broader patterns, with data from yeast indicating a dominant role in purifying selection in cis-regulatory evolution[20]. Allele-specific chromatin marks, particularly at imprinted gene loci, bring in added layers of [10]. Findings from Neanderthal introgressed alleles further underscore the dynamic nature of AI [21]. Beyond the realm of genetics, epigenetic factors, such as DNA methylation in sperm, interplay with AI. Parent-of-origin effects, influenced by aspects like parental age and BMI, impact a wide range of traits and diseases, deepening AI’s footprint [22–26]. While this inherent variance has been recognized in the realm of genetics for some time, its implications in complex disorders such as obesity remain underexplored.

Its significance extends through various tissues, developmental stages, and genetic studies, demonstrating the complexity and importance of AI [27–29]. Several genetic mechanisms, including parent-of-origin specific gene expression and genomic imprinting, underscore AI’s role in developmental potential and stability [30,31]. Moreover, AI provides insights into X-link gene dynamics, and the evolutionary trajectory of imprinted genes[10,32–34]. The realm of metabolic disorders, particularly obesity, captures the profound influence of AI, emphasizing its role in the intricate interaction between genetic predispositions and environmental factors [35,36], raising questions about the underlying genetic mechanisms and their implications for diseases such as obesity [37–39].

Our study situates itself amidst this vast landscape, seeking to demystify the role of specific classes of allelic imbalances in obesity and associated metabolic diseases based on a context-dependent colocalization approach. Are genes with low allelic imbalances indicative of stable expression levels across individuals, and what might this mean for metabolic health? Alternatively, what insights can we glean from genes with pronounced allelic imbalances with regulatory context? These are the major questions driving our research. The genetic risk associated with obesity which is an enormous disease burden [40], often located in noncoding genome regions, beckon in-depth exploration [41,42]. Many of these loci have been linked to adaptations of adipose tissue and hold implications for various metabolic diseases[35,43]. With adipose tissues taking center stage, the expanse of obesity’s influence is far-reaching, mediating metabolic diseases and contributing to systemic inflammation[35,36]. In conditions like type 2 diabetes mellitus (T2DM), an approach like colocalization unravels the roles of regulatory variants in disease susceptibility in immune cells in adipose tissues of obese individuals [44–46].

Through a comprehensive colocalization of the transcriptome, epigenome, and GWAS, we aim to decipher the intricate interplay of genetic variation, epigenetic regulation, and metabolism. This context-dependent colocalization approach helps us to identify common regulatory elements and pathways, revealing not just how genes, both protein-coding genes and long non-coding RNA (lncRNA)s, are regulated but also the mechanisms that underlie their dysregulation in pathological states like obesity. Our findings shed light on the genetic and epigenetic underpinnings of obesity and its associated susceptibility to other complex diseases.

## RESULTS

### Unraveling the Genetic Architecture of Obesity through Allelic Imbalance and Multi-omics Colocalization

To provide an intricate understanding of obesity, we utilized an array of multi-omics data sets (**Figure 1**). The transcriptome is primarily sourced from the Genotype-Tissue Expression (GTEx), containing bulk-tissue RNA-Seq data for about 838 samples across 49 tissues, including adipose tissues. We also leveraged epigenomic data consisting of DNase I hypersensitive sites (DHSs), with 3.6 million annotated sites across 438 cell and tissue types. Further, summary statistics of complex traits and disease associations were derived from GWAS catalogs, which included a total of all SNP-trait associations with genome-wide significance (p ≤ 5.0×10^-8^) from 6545 publications and millions of risk variants in complex traits, including body measurement, lipid or lipoprotein measurement, and metabolic disorder.

**Figure 1.**
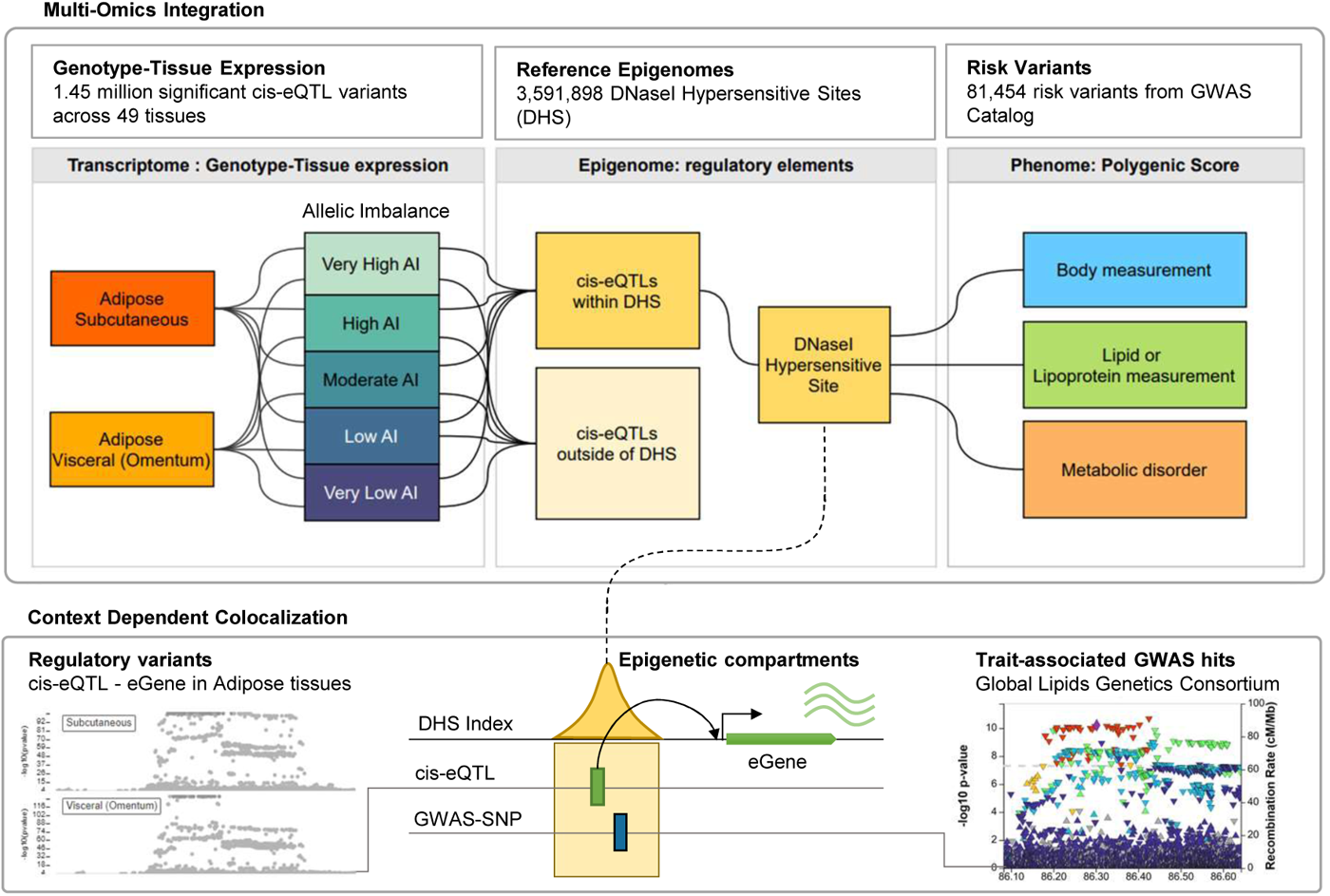
Study overview. This schematic illustrates the context-dependent colocalization analysis of allelic imbalance utilizing gene expression data from adipose tissues in obesity, combined with cis-eQTL, DNaseI hypersensitive sites (DHS), and GWAS variants.

Firstly, cis-expression quantitative trait loci (cis-eQTLs) offer a more focused and specific avenue for examining the regulatory effect of variants. Their inclusion in the analysis further refines our understanding of how genetic variants can affect gene expression. In the setting of DHS, cis-eQTLs assume an even more pronounced role. Here, they serve as a dual layer of refining the regulatory potential of eVariant-eGene pairs, affecting not only allelic imbalance but potentially influencing disease outcomes as well.

DNase I hypersensitive sites (DHS) serve as vital regulatory landmarks in the genomic landscape. Variants occurring within these sites have a heightened potential to affect gene regulation, either by altering transcription factor binding or by modifying other regulatory elements such as promoters, enhancers, silencers, and insulators. Given their importance, these adipose-associated cis-eQTLs within DHS become crucial for investigating the genetic regulation of obesity.

The analytical power of this study stems from its colocalization approach, integrating both allelic imbalance and regulatory variant data in the form of cis-eQTLs within DHS. This allows us to sift through the complexities of the regulatory landscape and pinpoint specific genes and regulatory variants that contribute to obesity. Moreover, this colocalization approach enables us to systemically discover novel candidate genes linked to regulatory variants, which include risk variants with regulatory effects as well as those target genes, including protein-coding gene and long non-coding RNA (lncRNA), not mapped based on LD-based approach in genome-wide association studies (GWAS). In doing so, we construct a comprehensive view of the myriad factors contributing to obesity, from the genetic to the epigenetic underlying complex diseases.

### Characterizing Allelic Imbalance Levels in Adipose Tissues and Their Pleiotropic Impact Across Tissues

We categorized allelic imbalance into five distinct levels: very low, low, moderate, high, and very high, based on the percentiles of allelic fold change (aFC) associated with 870,811 cis-eQTLs observed in adipose tissues (**Fig. 2A**). This stratification allowed us to precisely assess the differential impacts of allelic imbalances on gene expression and regulatory mechanisms. A total of 10,769 PCGs and 3,007 lncRNAs were identified and stratified into these groups based on their quantile aFC.

**Figure 2.**
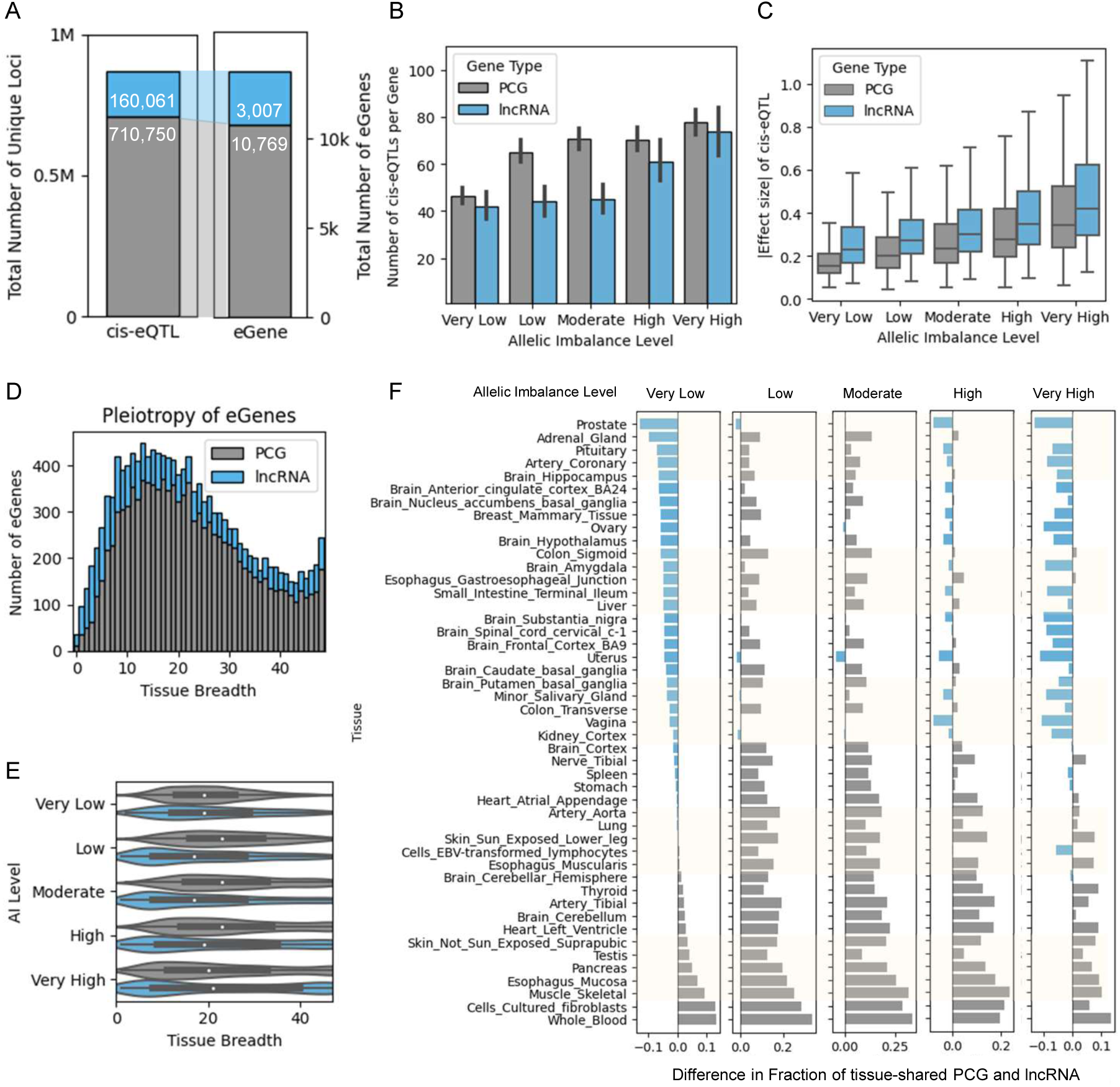
Tissue-specificity of cis-eQTLs between allelic imbalance levels. **A.** Total number of cis-eQTL and eGenes in adipose tissues. **B.** The mean absolute allelic fold change (aFC), respectively, the mean number of cis-eQTLs associated with adipose tissues per gene across different levels of allelic imbalance for protein-coding genes (PCGs) and long non-coding RNAs (lncRNAs). **C**. The distribution of effect sizes in adipose tissues for PCGs and lncRNAs. **D**. Distribution of the tissue-specific expression breadth of these genes across 49 tissues. **E.** Tissue breadth across different levels of allelic imbalance. **F.** Difference in fraction of shared genes between PCG and lncRNA at each level of allelic imbalance across tissue types.

Significant differences were observed in the average number of cis-eQTLs across AI levels (**Fig. 2B**). Specifically, for PCGs, the mean number of significant cis-eQTLs varied significantly across the different AI groups (P < 0.05). lncRNAs of very low AI and low AI groups were the only exceptions, where no significant differences in the number of cis-eQTLs were observed.

In adipose subcutaneous and adipose visceral tissues, we further dissected the genes and regulatory elements involved (**Fig. 2C**). In PCGs, the average absolute value of slopes differed 2.5-fold across AI levels, ranging from 0.1566 ±0.0001 in the very low group to 0.4352 ±0.0002 in the very high group. A similar trend was observed in lncRNAs.

Although AI levels were defined based on adipose tissue gene expression, we extended the analysis to a tissue breadth study across 47 tissues (**Fig. 2D); Additional data: Fig. S1**). In PCGs (**Fig. 2E**), genes with very low AI had significantly less tissue breadth than those with high (Mann-Whitney U test; p-value=0.032) and moderate (p=0.021) AI levels. For lncRNAs (**Fig. 2E**), genes with the highest AI level showed significantly larger numbers of shared genes than those at the lowest AI level (p=0.027).

Pleiotropic effects of cis-eQTLs appear to be widespread and show similar levels of allelic imbalance across different tissues in PCG rather than lncRNA, except lncRNAs with very low and very high AI levels (**Fig. 2F; Additional data:Fig. S2**). In PCGs, the average number of shared eGenes were 888.4 ± 77.0 (S.E.M) in the very low group and 1106.3 ± 65 in the high group, showing a significant difference (P = 0.032). In lncRNAs, the corresponding numbers were 259.3±19.4 and 310.5±11.4, also demonstrating significant difference (p=0.027).

The observed relationship between levels of allelic imbalance in adipose tissues and increased tissue sharing across multiple tissues captures intriguing difference in gene regulation and expression patterns. While allelic imbalance (AI) is captured through allele-specific expression (ASE) in adipose tissue, it’s crucial to clarify that a higher degree of AI implies potentially larger expression difference between individuals with different genotypes, and does not equate to tissue-specific expression. Genes with higher AI levels, as assessed in adipose tissues, were observed to have broader expression profiles across multiple tissues, pointing to a more pleiotropic role. This also suggests that genes with higher allelic imbalance could be players in biological processes that are across multiple tissue types, thus making them candidates for investigations into system-wide or multi-tissue disorders. Conversely, genes with low AI levels tend to have more tissue-restricted expression, indicating higher tissue specificity, indicating a more specialized role in adipose tissue functionality.

### Epigenetic Context-Based Refinement of Regulatory Potential of cis-eQTLs and eGenes

Our comprehensive analysis elucidates a clear correlation between the level of allelic imbalance and the density of cis-eQTLs within DNase I hypersensitive sites (DHS), as well as the genomic architecture of the corresponding genes (**Fig. 3A**). About 186,256 (26.2%) and 43,248 (27.%) cis-eQTLs fall into 154,475 and 36,720 DHSs, in protein-coding genes (PCGs) and long non-coding RNAs (lncRNAs), respectively (**Fig. 3B**). Only 1,671 DHSs are shared between PCG and lncRNA. In both PCGs and lncRNAs. The number of DHSs with cis-eQTLs per gene increased concomitantly with the level of allelic imbalance (**Fig. 3C**). Remarkably, the difference in the number of cis-eQTLs and DHSs between AI groups outweighed the difference observed between PCGs and lncRNAs, underscoring the significance of allelic imbalance as a determinant of regulatory complexity. Our statistical analyses further indicated a linear relationship between the number of DHSs and the number of cis-eQTLs per gene, substantiating the notion that allelic imbalance may serve as a proxy for assessing the regulatory landscape surrounding a gene. This observation also suggests that genes with lower AI levels tend to have a limited number of regulatory elements, raising questions about the properties of the genomic architecture of cis-eQTL within DHS and their target eGene.

**Figure 3.**
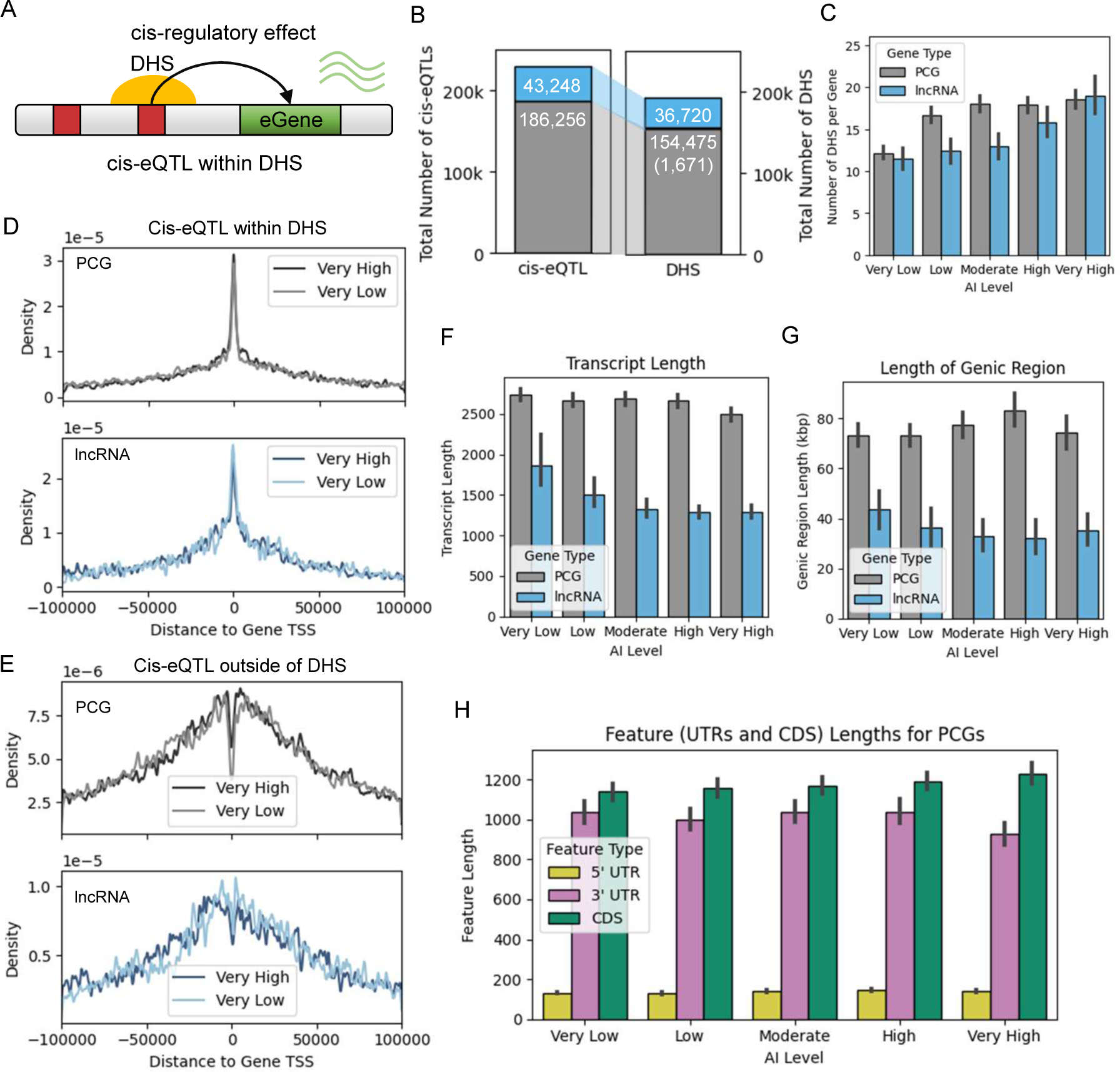
Characteristics of cis-eQTL located within DNase I Hypersensitive Sites (DHS) and their target genes. **A.** Schematic diagram of refining regulatory variants from cis-eQTL located with DHS. **B.** The count of cis-eQTLs located within DHS for both PCGs and lncRNAs. **C**. The average number of cis-eQTLs and DHSs linked to cis-eQTLs per gene across varying levels of allelic imbalance. (**D,E)** Distribution of cis-eQTL within DHS (**D**) and outside of DHS (**E**) around transcription start sites of target eGene. (F-H) The average transcript length (**F**), span of genic region (**G**) length (**F**), and lengths of gene features (**H**) within each AI category.

We show that cis-eQTL within DHS is located densely at the TSS (**Fig. 3D**), whereas those outside of DHS are distributed away from the TSS (**Fig. 3E**), implying a striking positional bias of cis-eQTLs, with those located within DNase I hypersensitive sites (DHS) being densely concentrated at transcription start sites (TSS). The proximity of these cis-eQTLs to TSS within accessible chromatin regions suggests a direct and potent influence on the transcriptional initiation of associated genes (**Fig. 3D**). This observation underscores the regulatory significance of cis-eQTLs in the genomic architecture of gene expression. The differential distribution of cis-eQTLs, with a subset situated away from TSS in non-DHS regions (**Fig. 3E**), further illuminates the nuanced landscape of genetic regulation. Such loci may represent distal regulatory elements or indicate indirect mechanisms of gene expression modulation. Our analysis encapsulates this spatial relationship, providing compelling evidence that genetic variants situated within DHS are more likely to exert functional effects by modulating transcription factor accessibility and chromatin dynamics.

Intriguingly, while higher levels of AI correlated with greater regulatory complexity, they were also associated with significantly shorter transcript lengths (**Fig. 3F**). For instance, genes at lower AI levels had mean transcript lengths of 2741.3±39.1 and 1829.2±153.6 for PCGs and lncRNAs, respectively. In contrast, these lengths considerably reduced to 246.0±39.1 and 1303.4±37.0 for both PCGs and lncRNAs at higher levels of AI (Mann-Whitney rank-sum test; P-value=6.7e-11 and P-value=1.5e-9). This inverse relationship between AI and transcript length could imply that genes with higher allelic imbalance are perhaps more finely regulated at the transcriptional level. Total genic regions in higher AI groups were also significantly larger than those in lower AI groups in PCGs (p-value=2.9e-25), but not in lncRNAs (**Fig. 3G**).

Diving deeper into the gene architecture, we discovered differences in untranslated regions (UTRs) across AI levels (**Fig. 3H**). The 5’ UTR was significantly longer in high AI groups (p-value=4.6e-6), while the 3’ UTR was considerably longer in lower AI groups (p-value=2.5e-37). Interestingly, the coding sequence (CDS) lengths showed no such divergence.

The differential between groups of varying allelic imbalance far outweighs the differences observed between PCGs and lncRNAs, accentuating the paramount influence of allelic imbalance on regulatory complexity. Moreover, statistical tests corroborate a linear association between the number of DHSs and cis-eQTLs per gene, underlining the potential of allelic imbalance as an informative metric for scrutinizing a gene’s regulatory landscape.

Our observations suggest that allelic imbalance serves as more than a mere indicator of differential gene expression between individuals; it is a robust gauge of a gene’s regulatory complexity and genomic structure. The convergence of higher allelic imbalance with increased regulatory involvement, paired with reduced transcript lengths, presents an intriguing avenue for future investigations. This inverse correlation suggests that genes with high allelic imbalance could be under more intricate transcriptional control. These findings could be especially relevant for exploring how genes with differing levels of allelic imbalance participate in complex biological networks and may contribute to disease susceptibility or phenotypic diversity.

### Functional Characterization of Allelic Imbalance in Adipose Tissues Across Metabolic Conditions

Differential gene expression in adipose tissues is a pivotal factor in understanding metabolic health, particularly in conditions like obesity, insulin resistance, and Type 2 Diabetes (T2D). In our study, we utilized data from eight different studies comparing gene expression profiles in adipose tissues from lean individuals and those who are obese but vary metabolically—from metabolically healthy to those with normal glucose tolerance, insulin resistance, and T2D (**Additional file: Table S1**). Differentially Expressed Genes (DEGs) were initially identified in each study (**Additional file: Fig. S3**) and then aggregated into overarching sets of up- and down-regulated genes. This comprehensive approach facilitated the exclusion of 47 genes that displayed contradictory patterns of expression across different studies. Overall, 2,725 up-regulated and 1,473 down-regulated genes were detected across various types of adipose tissues (**Fig. 4A**).

**Figure 4.**
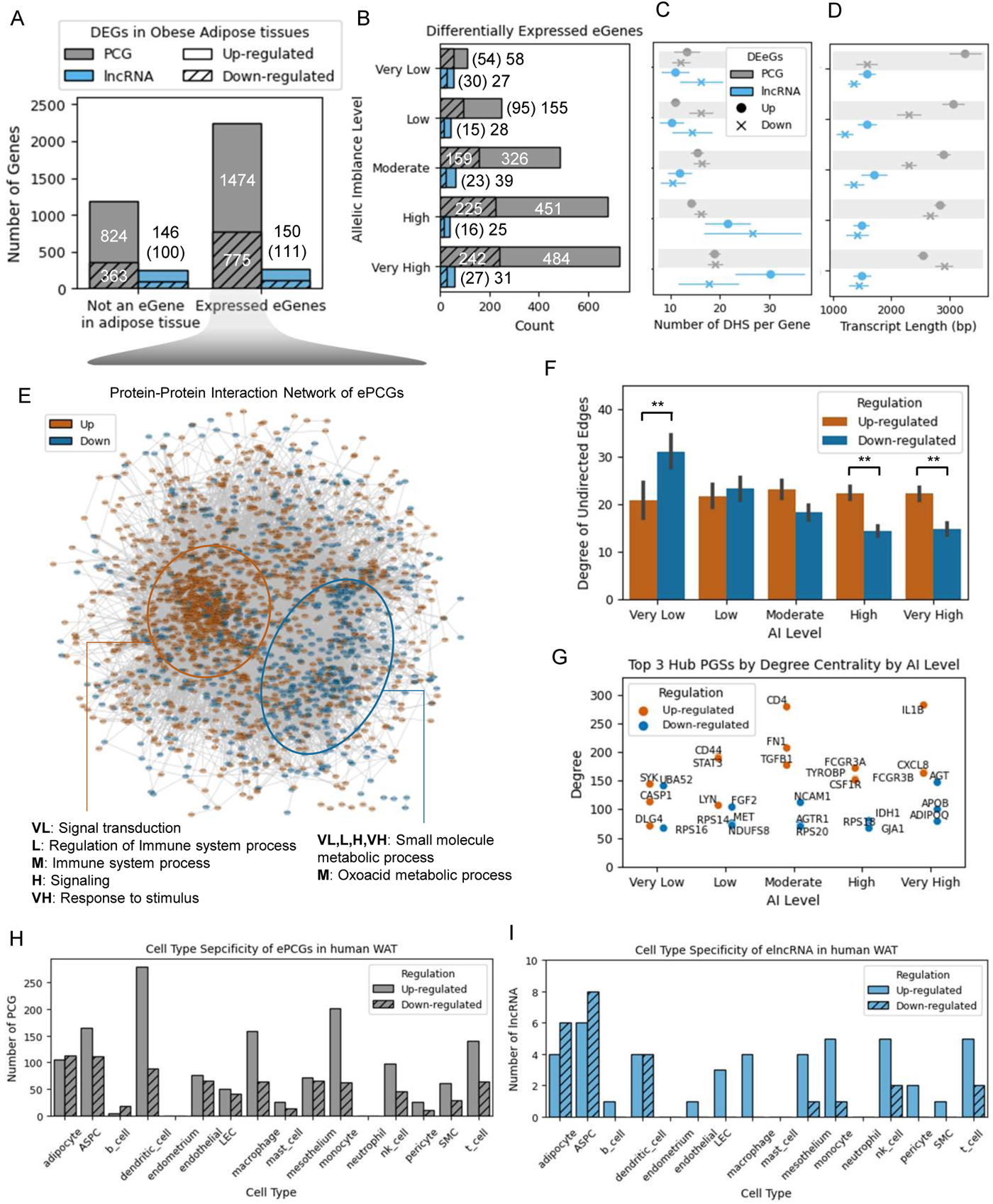
Differentially expressed eGenes in adipose tissues from individuals with obesity. **A**. Total number of differentially expressed PCGs and lncRNAs **B.** Number of differentially expressed eGenes across AI levels. **C,D**. The relationship between number of DHSs (**C**) and transcript length (**D**) in each AI level is depicted, with up-regulated PCGs having shorter transcripts and higher AI. Down-regulated PCGs having longer transcripts and higher AI. **E**. Protein-protein interaction network for the differentially expressed protein coding eGenes in obese adipose tissues. **F**. Degree of interacting proteins in DE protein-coding eGenes across AI levels. **G**. list the top 3 hub genes that are up- and down-regulated in obesity, respectively. **H.I** Cell type-specific expression heterogeneity of differentially expressed PCG (H) and lncRNA(I) in single cell analysis of human adipose tissues from individual with obesity.

As expected from the observation of the enrichment of regulatory elements in genes with higher AI levels, the up- regulation of genes was significantly more prevalent in moderate to very high levels of allelic imbalance (**Fig. 4B**). This trend was predominantly observed in protein-coding genes and was remarkably absent in long non-coding RNAs (lncRNAs). These findings point toward the potential functional ramifications of allelic imbalance in metabolic health, particularly its potential role in exacerbating conditions like obesity and insulin resistance.

DNase I hypersensitive sites (DHS) density—a marker for chromatin accessibility—also displayed AI-level-dependent variation (**Fig. 4C**). For both up- and down-regulated genes, DHS density was significantly higher in the higher AI group than in the lower AI group (p=0.003 and p=0.015, respectively). Moreover, up-regulated long non-coding RNAs (lncRNAs) manifested a higher DHS density in the highest AI group, compared to their counterparts in lower AI groups (Figure S6, p=0.007). These observations offer compelling evidence that chromatin structure and accessibility could serve as vital mechanisms underpinning gene regulation across different AI strata.

Our meticulous examination of transcript lengths in protein-coding genes (PCGs) unearthed intriguing disparities that could be functionally impactful (**Fig.4D**). Specifically, we observed that up-regulated genes within lower AI groups possess significantly larger transcript lengths compared to those in higher AI groups (p=0.010). Conversely, down-regulated genes in higher AI groups exhibited substantially larger transcript lengths than those in lower AI groups (p=3.4e-9). These disparate findings not only underscore the complexity of gene regulation but also suggest that transcript length could be a discriminating feature in the regulation of gene expression across different AI levels.

Protein-protein interaction (PPI) networks further substantiated the functional characterization of these differentially expressed protein coding genes (**Fig. 4E**). Here, a clear dichotomy was observed, wherein one cluster primarily comprised up-regulated genes, while another cluster contained mostly down-regulated genes. Our functional enrichment analysis reveals stark differences in biological processes influenced by both up- and down-regulated genes across various AI levels (**Additional file: Fig. S4**). Critically, the up-regulated genes in very high AI groups were statistically enriched (p-value=0.0024) for roles in ‘response to stimulus.’ Conversely, those in the very low AI groups were linked to ‘Signal Transduction’ (Figure 6A). This functional divergence was further nuanced for genes in the low AI, moderate, and no cis-eQTL groups, which were enriched in ‘Immune System Process.’ Down-regulated genes presented another layer of complexity. Except for the moderate AI group, these genes prominently figured in ‘Small Molecule Metabolic Process’ (Figure 6B). In the very low and low AI groups, as well as the no cis-eQTL group, there was significant involvement in ‘cytoplasmic translation’ (p-value=0.0007).

**Figure 5.**
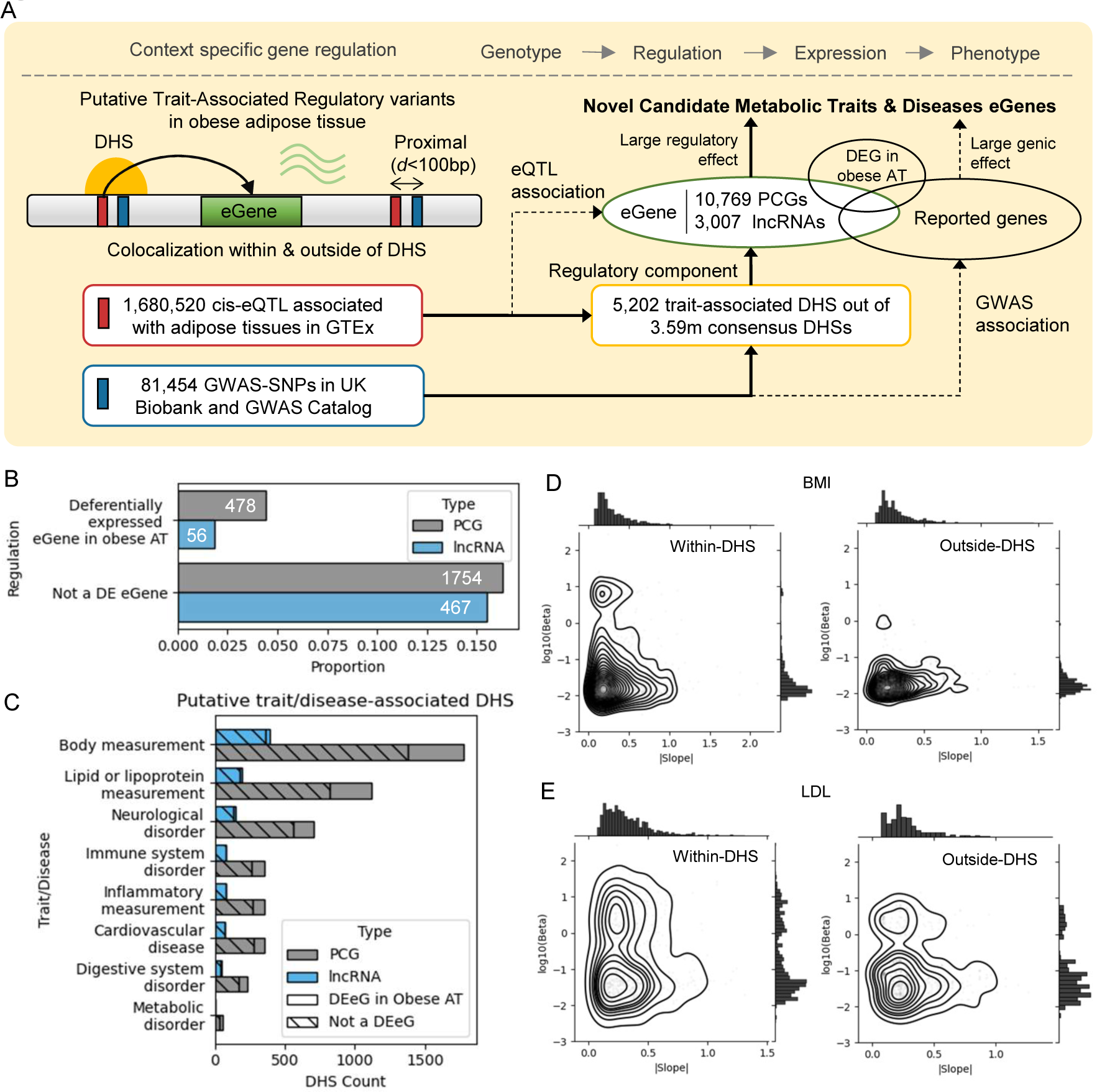
Context-Dependent Colocalization of cis-eQTL and GWAS-SNPs within DHS. **A.** Schematic diagram of colocalization of cis-eQTL and GWAS-SNPs with DHS. GWAS-associated SNPs present within DHSs. **B.** Proportion of eGenes that are associated with cis-eQTL and GWAS-SNP within DHS for both PCGs and lncRNAs. **C.** Number DHS colocalized with cis-eQTL and GWAS-SNPs across complex traits and diseases. D.E. Conditional density of effect size (β) of GWAS-SNP given absolute slope value of cis-eQTL colocalized with DHS in BMI (**D**) and LDL (**E**).

**Figure 6.**
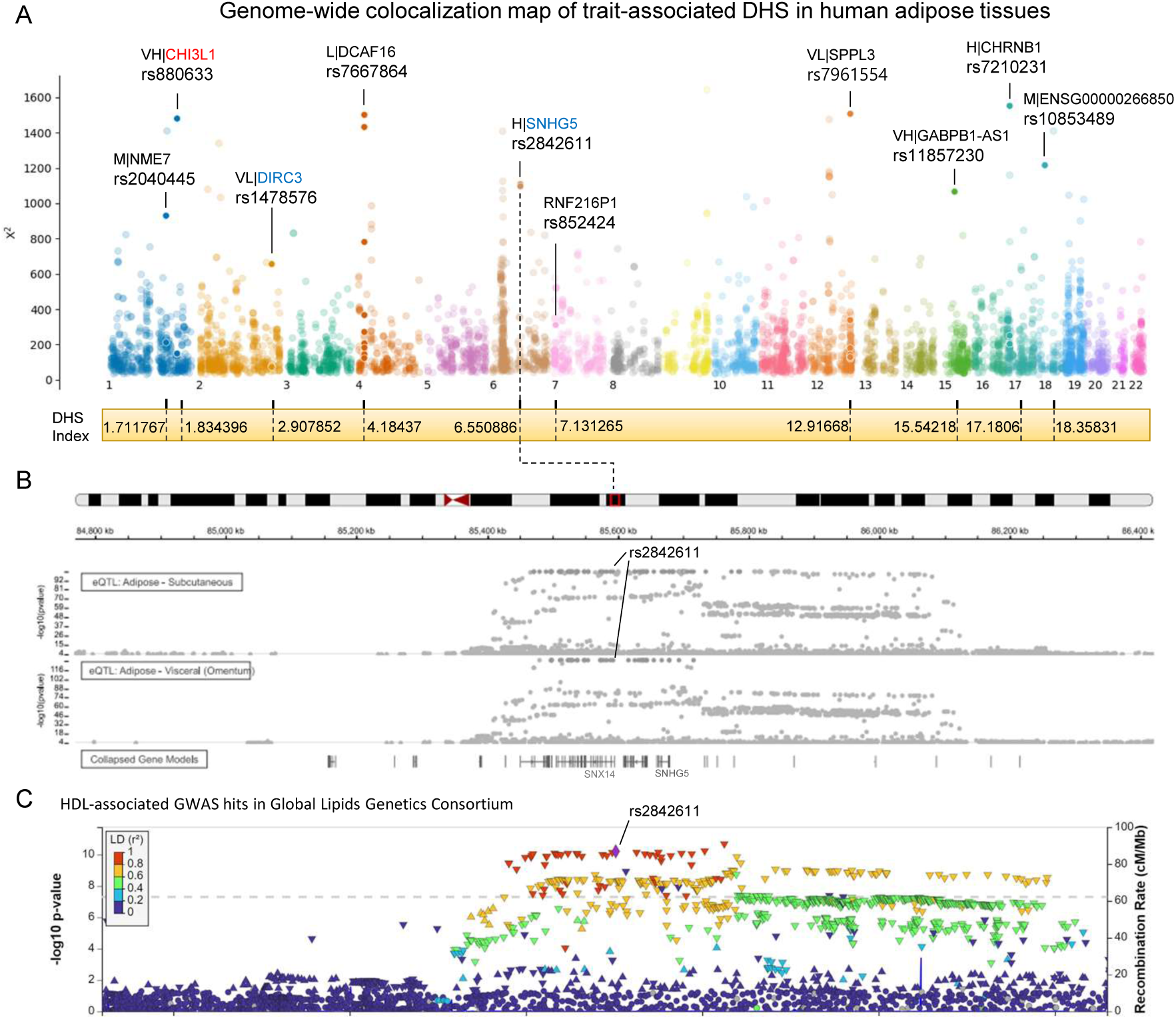
Genome-wide map of composite score of colocalization of cis-eQTL and GWAS-SNP within context of epigenic element. **A**.Genome-wide colocalization map of regulatory variants and trait-associated variants within context of epigenetic element based on composite probability. X is computed by using the Fisher’s method for composite probabilities of pair of cis-eQTL and GWAS-SNP within DHS. Gene name with red and blue depicts up-regulated and down-regulated in obese adipose tissues, respectively. A example of colocalization for SNHG5 linked to a DHS that have a variant associated with both cis-eQTL’s regulatory effect (**B**) and genic effect on complex traits (**C**).

Intriguingly, DEGs in the category of very high AI demonstrated a significantly lower degree of interacting proteins in these PPI networks (**Fig. 4F**), implying that these genes might play isolated yet potentially crucial roles within biological networks. The functional significance of these DEGs was also evident when focusing on hub genes—genes that interact with multiple other genes in the network (**Fig. 4G**). The preponderance of up-regulated genes in higher AI levels may indicate that allelic imbalance contributes to the severity of metabolic conditions. Notably, up-regulated genes in higher AI levels, such as CD4 and IL1B, are known to regulate immune responses, pointing toward their role in systemic inflammation commonly associated with obesity and metabolic disorders like insulin resistance. The distinction in hub gene expression, especially the up-regulation of CD4, IL1B, and FCGR3A in adipose tissues with obesity, provides a crucial insight into the underlying metabolic health. CD4 and IL1B are integral components of immune response and cellular communication, suggesting their heightened role in inflamed adipose tissue. Conversely, down-regulated hub genes like PPARG and APOE have established roles in lipid metabolism, indicating potential defects in adipocyte function and lipid storage in conditions of high AI. Additionally, down-regulated genes also showed a higher propensity to be hub genes, hinting at their critical role in the cellular network and metabolic pathways.

Finally, Single-cell expression data adds another layer of complexity, particularly pointing toward cellular heterogeneity in adipose tissues. The single-cell expression levels of these differentially expressed PCGs (**Fig. 4H**) and lncRNA (**Fig. 4I**) were investigated in human white adipose tissues, where the number of down-regulated genes is larger than those of up-regulated genes especially in adipocytes. Notably, up-regulated genes like CD4, PTPRC, and ITGAM were predominantly expressed in immune cells such as dendritic cells, macrophages, and monocytes (**Additional file: Fig. S5**). On the other hand, down-regulated genes like PPARG, APOE, and PTEN were more commonly expressed in adipose precursor cells and adipocytes.

The distinct clustering and centrality metrics observed in the PPI networks for DEGs across different AI levels suggest that genes with higher AI may serve as isolated but critical nodes within these networks, potentially functioning as key modulatory points in metabolic health. The identification of cell type-specific hub genes opens up avenues for therapeutic targeting, given their well-documented roles in metabolic pathways. Their down-regulation in high AI levels also emphasizes the need for targeted interventions in metabolic conditions like obesity and T2D.

### Conditional Analysis of Risk Susceptibility from Colocalization of cis-eQTLs and GWAS-Associated SNPs with regulatory context

A fundamental aspect of unraveling gene regulation in complex traits involves bridging the gap between cis-regulatory effects and established genetic risk of regulatory variants. Our method leverages the colocalization of DNase I hypersensitive sites (DHSs) with an extensive dataset of 1.6 million cis-eQTLs identified in adipose tissue and 81,454 SNPs from GWAS Catalog (**Fig. 5A**). This approach enables the identification of target genes for regulatory variants associated with risk susceptibility, encompassing both protein-coding genes (PCGs) and long non-coding RNAs (lncRNAs). Uniquely, our colocalization strategy excels in pinpointing GWAS-associated risk variants and elucidating their gene targets, going beyond conventional gene mapping based on linkage disequilibrium coordinates. This is particularly valuable for regulatory variants located in non-coding regions, such as introns of proximal genes, where traditional methods may overlook functional connections.

In our analysis, we identified 5,202 DHSs that exhibit colocalization with cis-eQTLs and GWAS-SNPs, revealing a significant enrichment of these DHSs among differentially expressed protein-coding genes (PCGs) across a range of complex traits and diseases (p-value < 0.002, Fig. 5B). Within this subset, 478 (21.4%) PCGs and 56 (10.7%) lncRNAs differentially expressed in obese adipose tissues were linked to variants that demonstrate both regulatory and genic effects. This notable intersection of gene expression dysregulation in obesity with genetic susceptibility loci underscores the potential of these DHS-associated genes as candidate mediators of obesity’s molecular mechanisms.

These findings affirm the complex interplay inherent to cis-regulatory effects, illustrating their contingent influence on the genetic predisposition to obesity and related traits, such as body measurements and lipid or lipoprotein levels (**Fig. 5C**). Furthermore, the observed colocalization between GWAS-associated SNPs and DHSs sheds light on the role of chromatin accessibility in shaping these intricate genetic relationships.

To delineate the interplay of genetic effects for cis-eQTL and GWAS-SNP pairs colocalized within DHS, we constructed a conditional density for a control subset comprising cis-eQTLs proximal to GWAS-SNPs but not colocalized with DHS, focusing on BMI (**Fig. 5D**) and LDL levels (**Fig. 5E**). Our analysis revealed that within DHS, the effect sizes(β) of GWAS-SNPs are conditionally amplified in the presence of cis-eQTLs with lower effect weights. In contrast, when paired proximally outside of DHS, this conditional enhancement of GWAS-SNP effect sizes is markedly diminished. These observations suggest that the regulatory landscape defined by DHS plays a pivotal role in modulating the genetic impact on obesity and lipid traits, potentially guiding more refined genetic risk assessments and therapeutic interventions.

We devised a composite probability of taking a sum of the natural logarithm of probabilities for cis-eQTL and GWAS-SNP colocalized within DHS. Since those cis-eQTL and GWAS-SNP are significant from independent tests from studies they are derived may be of very different, it is plausible to take into account only those probabilities to obtain a single test of the significance based on the product of the probabilities from individual studies, especially in a case where effect size of cis-eQTL and GWAS-SNP have asymmetrically related [47] as they are systematically different in functional annotation, selective pressure, and regulatory landscape [48]. We build a genome-wide colocalization map to novel genetic and epigenetic factors and prioritize them based on the composite scores based on GWAS and cis-eQTL studies (**Fig. 6A**) that follows Chi-square distribution of *n* = 2 (**Additional file: Fig. S6**). Out of those 5202 DHSs (**Additional file: Table S2**), we highlight a lncRNA SNHG5 to a top candidate from gene set with high allelic imbalance level from cis-eQTL analysis (**Fig. 6B**). A HDL-associated SNP, rs2842611, is located in an intron of the neighboring gene, SNX15, from GWAS (**Fig. 6C**), but, its cis-regulatory effect is linked to the expression level of SNHG5. This illustrates that the conventional gene mapping approach in GWAS based on chromosome coordinates may miss a complex regulation mechanism involved with SNHG5 that is down-regulated in obese adipose tissue. Our context-dependent colocalization approach identifies novel candidates with regulatory potentials of trait-associated variants and well as refines a gene set associated underlying obesity.

## DISCUSSION

In a world grappling with rising obesity rates, understanding the genetic factors behind this condition becomes ever more crucial. Our study sheds light on such genetic and epigenetic intricacies, particularly emphasizing the role of regulatory context with allelic imbalance in gene expression patterns associated with complex traits. By delving deep into both coding and non-coding genes, we’ve unveiled new insights about genetic predispositions. These findings not only enhance our comprehension of obesity’s genetic basis but also point to potential avenues for future research and therapeutic strategies.

By discerning these genetic risk susceptibility sites in AI’s context, we’re introducing an added dimension of complexity. While recent strides have been made in fusing GWAS outcomes with other genetic indicators, our study distinguishes itself by integrating these findings with regulatory context. This integrative approach not only deepens our grasp of obesity’s genetic risk landscape but also paves the way for similar explorations into other complex traits. Our colocalization approach bridges allelic imbalance (AI) with key genetic risk markers such as SNPs and DNase I hypersensitive sites (DHSs). Most of the lncRNA identified in this study are unexplored in their genetic mechanism in metabolic traits. Further experiment is needed to validate genetic mechanism of how regulatory variants of lncRNAs in obesity are associated with complex disease. Colocalization of these genetic risk variants with regulatory potential promises to fine-tune the accuracy of existing genetic risk models. Such refined models could substantially guide clinical choices, particularly concerning prevention and management strategies for conditions tied to obesity.

Our study identifies a quantitative pattern of interplay: higher AI levels correlate with up-regulated short genes accompanied by a larger number of DHSs, whereas their lower counterparts align with down-regulated long genes, but with fewer DHSs. The broader ramifications of these findings are manifold. Understanding this nuanced interplay between AI, gene length, and DHS abundance can open avenues for more individualized therapeutic interventions. For instance, such a well-known protein-coding gene, GLP1R, one of major drug targets for diabetes and obesity[49], is identified to have high allelic imbalance in cis-regulatory variants and associated with type 2 diabetes, which implies potentially large difference in expression level between individuals with reference allele and risk alleles. This precision might manifest in targeted treatments using pharmacological agents or advanced gene editing technologies at single cell. Furthermore, these AI profiles, combined with GWAS and DHS data, can act as potential biomarkers. Such markers can be instrumental in identifying individuals predisposed to obesity-linked complications, thereby bolstering preventive measures to be both timely and impactful.

Diverging from traditional differential gene expression explorations, our study delves deep into the functional richness and interactions among the DEGs. We chart novel territory by establishing the interplay between transcript length, UTR dimensions, and differential gene expression across diverse AI levels in obesity. Of particular note are our findings on the enrichment of immune response and cellular signaling pathways among up-regulated genes and metabolism in down-regulated genes, reinforcing their significance in disease pathways. Our findings not only underscore the importance of gene features in obesity but also highlight their connection with allelic imbalance and complex trait association. This offers a holistic view of obesity’s transcriptomic framework underlying metabolic trait.

Our study, while providing a comprehensive perspective on the nexus between regulatory effect and genic effect in epigenetic context associated with obesity, isn’t without its constraints. First, the treasure trove of high-throughput genomic data, while offering a depth of knowledge, is not without pitfalls. The challenges of data sparsity and embedded cryptic noise present hurdles that could sway the fidelity of our results. Second, the breadth of our investigation, though expansive on the genetic facets of obesity, falls short in robustly addressing the intertwined environmental influences central to this condition. The complexity of obesity arises not only from genetic underpinnings but also from myriad environmental factors such as diet, lifestyle, neurological intricacies, and societal health determinants. Third, drawing from trait-associated SNPs in large-scale GWAS, we unearthed invaluable insights into risk-associated SNPs with regulatory potential. However, the nuances of GWAS can sometimes tether its findings to particular population specifics, potentially sidelining the richness of genetic diversity inherent in mixed or heterogeneous populations. Lastly, our focus, honed in on specific tissues and cell types, may overlook the broader tapestry of tissues instrumental in obesity’s genesis and progression. This selectiveness inadvertently omits the nuanced dynamics of cell type heterogeneity inherent within these tissues. Such an oversight might lead to potential blind spots, obfuscating essential cell-specific influences on obesity.

Our acknowledgment of these limitations underscores our commitment to clarity and transparency. It’s a clarion call for future endeavors in this realm to bridge these gaps, ensuring a more integrated, comprehensive model of obesity that seamlessly melds genetic insights with environmental realities. The associations and non-coding genes spotlighted demand rigorous experimental validation to underscore their true functional relevance in the obesity landscape. Looking forward, the insights gained not only set a robust foundation for subsequent research endeavors but also spotlight potential avenues for improving diagnostic and therapeutic methodologies. Embracing this multidimensional approach, we envisage a future where precise diagnostics, proactive alert systems, and bespoke interventions become the standard for managing obesity and related complexities. It’s our hope that this study acts as a beacon, galvanizing interdisciplinary alliances aimed at enhancing individualized care for those at heightened risk of obesity-associated challenges.

## ACKNOWLEDGEMENTS

This work is supported by the Medical Research Center Program (2022R1A5A2018865) through the National Research Foundation of Korea (NRF) funded by the Korean Ministry of Science and ICT (MSIT).

## AUTHOR CONTRIBUTIONS

S.M designed the study with S.-Y.P. S.M run the experiments. All authors contributed to the development of the project. All authors participated in construction of the manuscript.

## METHODS

### Datasets Collection

#### Patient Samples and Gene Expression Data

We sourced RNA-sequencing data of human adipose tissues from both obese and lean individuals from the Gene Expression Omnibus (GEO). Specific GEO study IDs included are GSE179455, GSE165932, GSE55008, GSE156906, GSE205668, GSE110729, GSE141432, and GSE162653. We identified differentially expressed genes (DEGs) across various adipose tissue types, such as epicardial, omental, and subcutaneous tissues (Supplementary Table 1). Groups were classified based on obesity with/without insulin resistance, normal glucose tolerance (NGT), Type 2 Diabetes (T2D), and metabolically unhealthy conditions compared to lean metabolically healthy controls. A significance level cutoff of 0.05 was employed following Benjamini & Hochberg adjustment for P-values, with a fold change threshold of 1.5 (DESeq2, Love et al., 2014). DEGs that showed contradictory regulation across studies were excluded from further analysis.

#### Significant single-tissue cis-eQTLs

Cis-expression Quantitative Trait Loci (eQTL) data was acquired from the GTEx project V8, which comprises RNA-seq and whole-genome sequencing data from 49 tissues of 838 healthy individuals (dbGaP: phs000424.v8.p2). We extracted cis-eQTL variants below the predefined significance threshold and calculated the slope of significant eGenes using a q-value < 0.05, derived from beta-approximated permutation p-values.

#### Allelic Imbalance Classification and Tissue Breadth

cis-eQTL effect sizes were quantified as the allelic fold change in gene expression between reference and alternative alleles. We categorized the magnitude of allelic imbalance into five classes: ‘Very Low’, ‘Low’, ‘Moderate’, ‘High’, and ‘Very High’, based on {0, 20, …, 80, 100} percentile distributions of absolute log2-transformed allelic fold change (aFC) values in adipose tissues. For genes with multiple adipose tissue estimates, the largest absolute log2(aFC) was selected. Tissue breadth was assessed by the number of cis-eQTL target genes across the 49 GTEx tissue types.

#### Regulatory DNA Mapping and Genetic Association

In our approach, we employed a direct colocalization strategy to link SNPs identified in GWAS studies with regulatory DNA regions characterized by DNase I hypersensitive sites (DHSs). This analysis was designed to pinpoint potential regulatory SNPs that may contribute to complex traits. By overlaying GWAS SNPs onto the DHS Index data, we identified SNPs that fall within regulatory regions, potentially influencing gene expression. Additionally, we integrated this data with cis-eQTLs associated with genes present in these regulatory DNA regions. This colocalization enables the direct identification of candidate target genes that are not only implicated in trait variation but are also likely to be directly regulated by these SNPs. This streamlined approach underscores the potential regulatory mechanisms by which genetic variation can influence complex traits and provides a clear path to deciphering gene-trait association

#### Integration of Effect Sizes using Fisher’s Method

To assess the combined significance of colocalized pairs of cis-eQTLs and GWAS-SNPs within DHS, we employed Fisher’s method, a statistical approach that combines multiple independent p-values into a single test statistic. This class method is particularly suited for our analysis, as it allows us to integrate evidence from distinct genetic studies to evaluate the overall significance of observed associations [47].

Given m independent tests with p-values *p*_1_, *p*_2_, …,*p*_m_, Fisher’s method combines these into a single test statistic, *X*^2^, defined as:

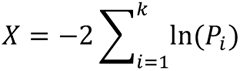

Under the null hypothesis that all the individual tests are independent, *X*^2^ follows a chi-square distribution with 2*m* degrees of freedom. We then calculate the composite p-value from this chi-square distribution to assess the joint significance of the colocalized variants within DHS.

In the context of our study, *m*=2 for each pair of colocalized cis-eQTL and GWAS-SNP within DHS, reflecting the integration of genetic effects from both regulatory and trait-associated perspectives. We constructed conditional densities for subsets of colocalized pairs both within and outside DHS to compare the interaction of their genetic effects. We constructed null model of context-dependency in colocalization between cis-eQTL and GWAS-SNP by obtaining pairs of cis-eQTL and GWAS-SNP located within distance less than 100bp apart. The conditional probability of observing a GWAS-SNP’s effect size, given the effect size of a proximal cis-eQTL, was evaluated to discern patterns of genetic influence modulated by the presence of DHS. We identify subset of candidates conditional given allelic imbalance level of cis-eQTL since there can be a discovery tradeoff between regulatory effect and genic effect on the gene[48].

This analysis enabled us to identify genetic loci where the combined evidence from cis-eQTLs and GWAS-SNPs suggests a significant regulatory impact on complex traits, underscoring the utility of Fisher’s method in uncovering potential genetic mechanisms underlying trait variation

#### Protein-Protein Interaction (PPI) Network Analysis

PPI networks were generated using high-confidence interactions from the STRING database, considering only interactions with a confidence score above 0.7. Network construction and analysis were performed using Cytoscape, integrating differentially expressed genes (DEGs) with varying levels of allelic imbalance. Topologically significant nodes, characterized by high connectivity, were identified by using Network Analyzer. These key hub genes were then used to reconstruct the network, highlighting potential pathways and biological processes that might be disrupted in obesity and associated metabolic disorders.

#### Single cell expression heterogeneity analysis

Single cell RNA-seq data of human white adipose tissue from individuals with obesity [50] were analyzed to assess the cellular distribution of DEGs associated with allelic imbalance. Expression profiles were visualized using 2D Uniform Manifold Approximation and Projection (UMAP) plots.

#### Functional Enrichment Analysis

Enrichment analyses for biological pathways were conducted using g:Profiler with differentially expressed protein coding genes (PCG) across allelic imbalance levels. Biological processes and pathways with a false discovery rate (FDR) below 0.05 were reported as significantly enriched.

#### Statistical analysis

All statistical tests employed are specified in corresponding text and figure legends. For non-parametric distributions, the Mann-Whitney U test was utilized. The hypergeometric test was applied for evaluating the overlap between gene sets. Analyses were conducted using appropriate Python and R packages, with multiple hypothesis correction applied using the Benjamini-Hochberg procedure where appropriate.

#### Gene annotation

Gene features such as biotype, UTR lengths, and transcript lengths were extracted from the ENSEMBL BioMart database. Protein-coding genes were defined based on the presence of an associated protein product, excluding transcripts classified as retained introns or those undergoing nonsense-mediated decay. Long non-coding RNAs (lncRNAs) were identified based on the biotype provided in the ENSEMBL annotations.

## Data Availability

The datasets and resources used in our study are publicly accessible via the provided URLs. The GTEx V8 cis-eQTL and eGene data are available at the GTEx portal. DHS Index data can be accessed through the dedicated repository.

The GWAS Catalog, and ENSEMBL BioMart provide extensive databases for genetic, epigenetic, and gene annotation resources. All data used for analyses are available within these resources.

GTEx Analysis V8 cis-eQTL and eGene: https://storage.cloud.google.com/adult-gtex/bulk-qtl/v8/single-tissue-cis-qtl/GTEx_Analysis_v8_eQTL.tar

DHS Index: https://www.meuleman.org/DHS_Index_and_Vocabulary_hg38_WM20190703.txt.gz

A single-cell atlas of human white adipose tissue: https://singlecell.broadinstitute.org/single_cell/study/SCP1376/a-single-cell-atlas-of-human-and-mouse-white-adipose-tissue?cluster=Human%20WAT&spatialGroups=--&annotation=cluster--group--study&subsample=all#study-visualize

Imprinted gene database: https://www.geneimprint.com/site/genes-by-species.Homo+sapiens GWAS Catalog: https://www.ebi.ac.uk/gwas/api/search/downloads/alternative

Ensembl BioMart: https://useast.ensembl.org/biomart/martview/

## Code Availability

The complete set of scripts and codes used for data analysis and visualization in this study are maintained and documented on GitHub at: https://github.com/moon-s/allelic-imblance

## Supplementary tables

**Table S1.** RNA-Seq of adipose tissues from individuals with obesity and metabolic disease states.

| GEO ID | Control group | Test group | Tissue/cell |
| --- | --- | --- | --- |
| GSE179455 | Lean | Obese/overweight | Epicardial adipose tissue |
| GSE165932 | Lean (IS) | Obese (IR) | Omental adipose tissue |
| GSE55008 | Lean | Obese | Omental adipose tissue |
| GSE156906 | Lean | Obese (MUH) | Subcutaneous adipose tissue |
| GSE156906 | Lean | Obese (MH) | Subcutaneous adipose tissue |
| GSE205668 | Normal weight | Obese | Subcutaneous adipose tissue |
| GSE110729 | Lean | Obese | Subcutaneous white adipose tissue |
| GSE141432 | Lean | Obese (NGT) | White adipose tissue |
| GSE141432 | Lean | Obese (T2D) | White adipose tissue |
| GSE162653 | Lean | Obese | White adipose tissue |

\*IS: insulin sensitive, IR: insulin resistance, MUH: metabolically unhealthy, MH: metabolically healthy, NGT, Normal glucose tolerance, T2D: type 2 diabetes

Table S2. Composite probability of colocalization between cis-eQTL and GWAS-SNP based on Fisher’s methods

## Supplementary figures

**Figure S1.**
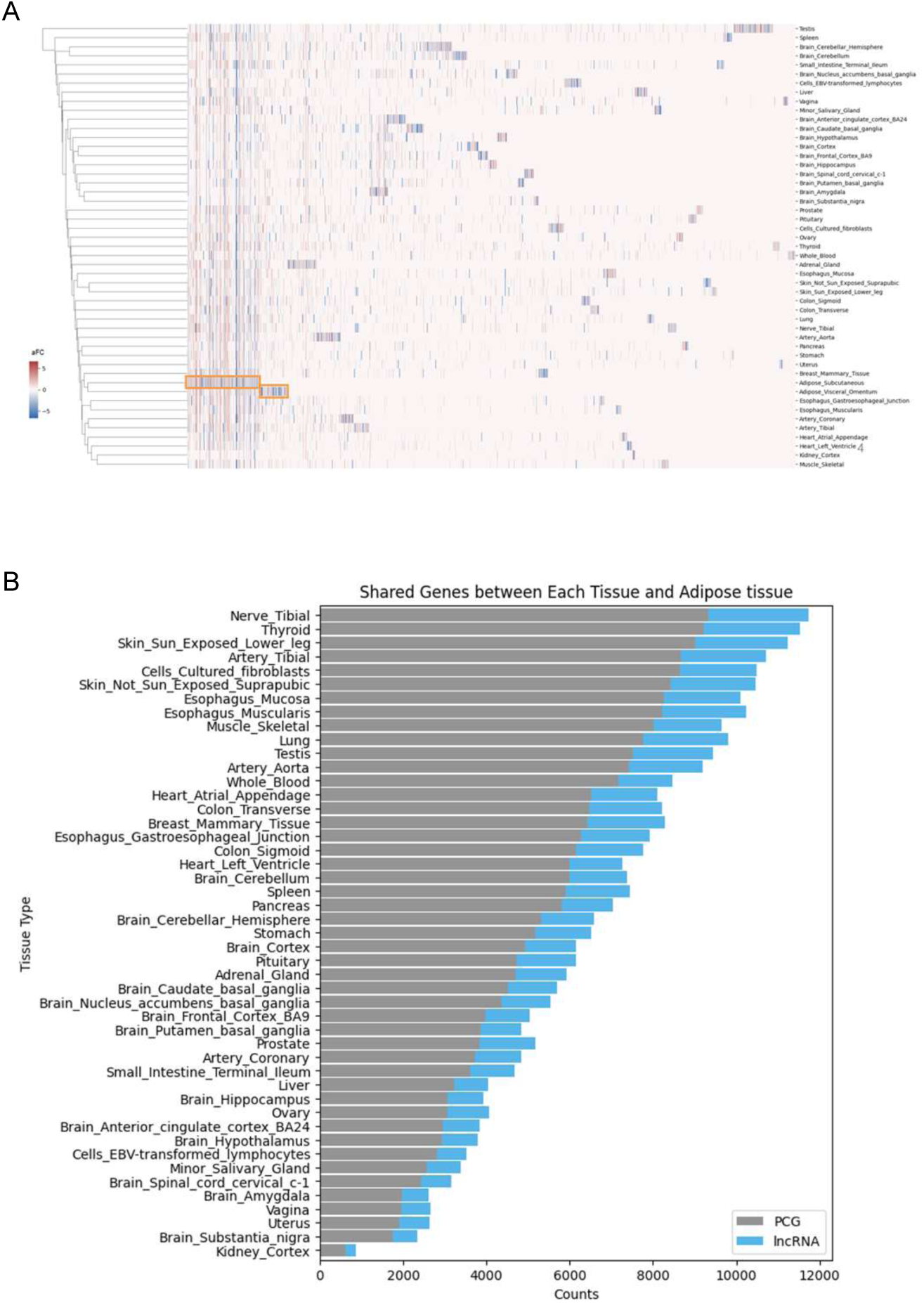
Tissue-Specificity of eGenes in adipose tissues. **A**.the heatmap of the allelic fold change (aFC) of genes with significant cis-eQTLs in each tissue types. The x-axis represents individual genes, and the y-axis represents tissue types. A distinct cluster of eGenes in adipose tissues is highlighted in orange, suggesting small fraction of genes show tissue-specific regulatory mechanisms. **B.** Number of eGenes shared between adipose tissues, subcutaneous and/or visceral omentum, and other tissues types.

**Figure S2.**
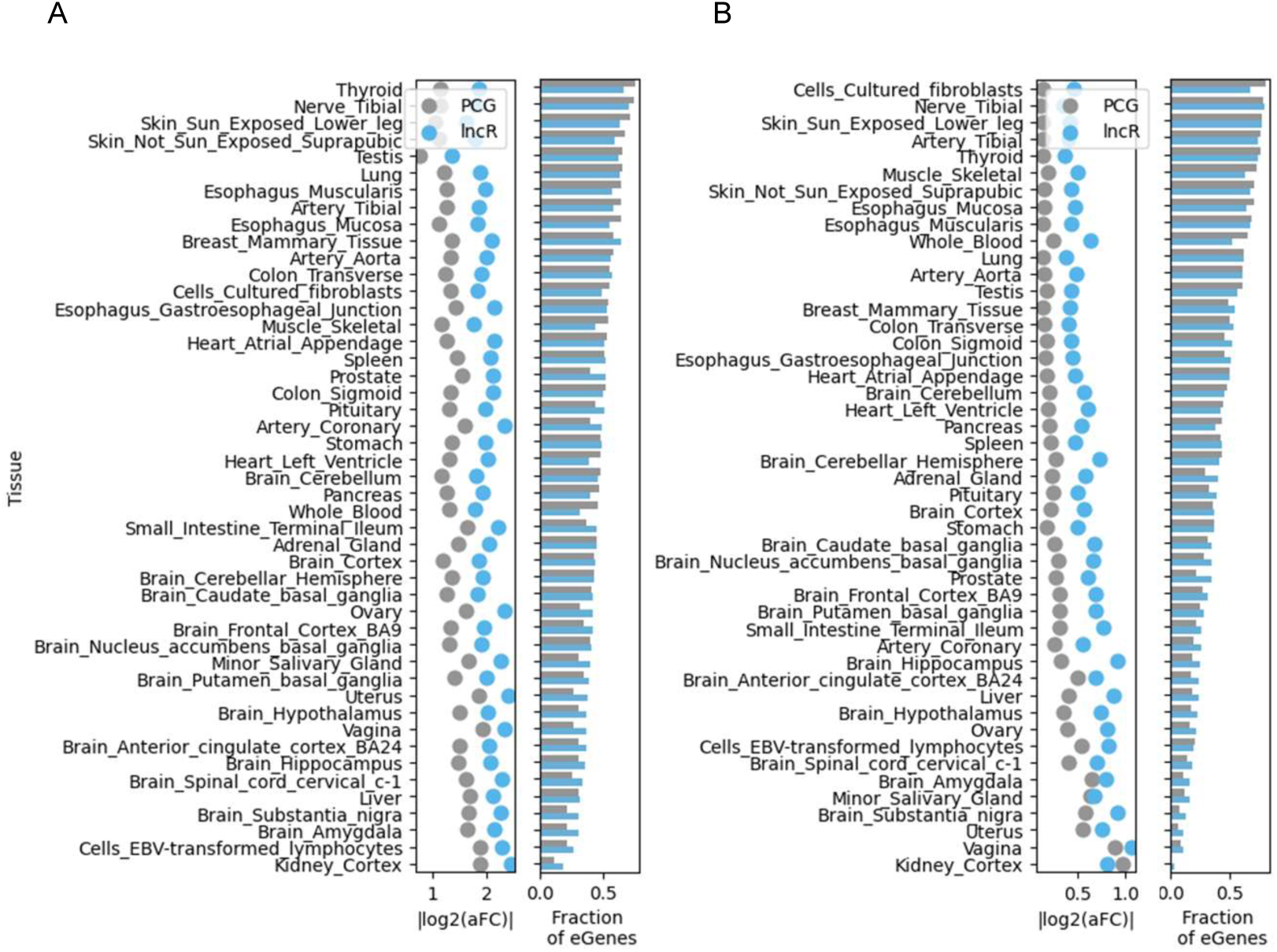
Allelic fold change of shared eGenes across tissue types. For shared eGenes between adipose tissue and other tissue, we computed fraction of eGenes shared between adipose tissue and each tissue, and mean aFC of them in each tissue for very high allele imbalance (A) and very low allele imbalance (B)

**Figure S3.**
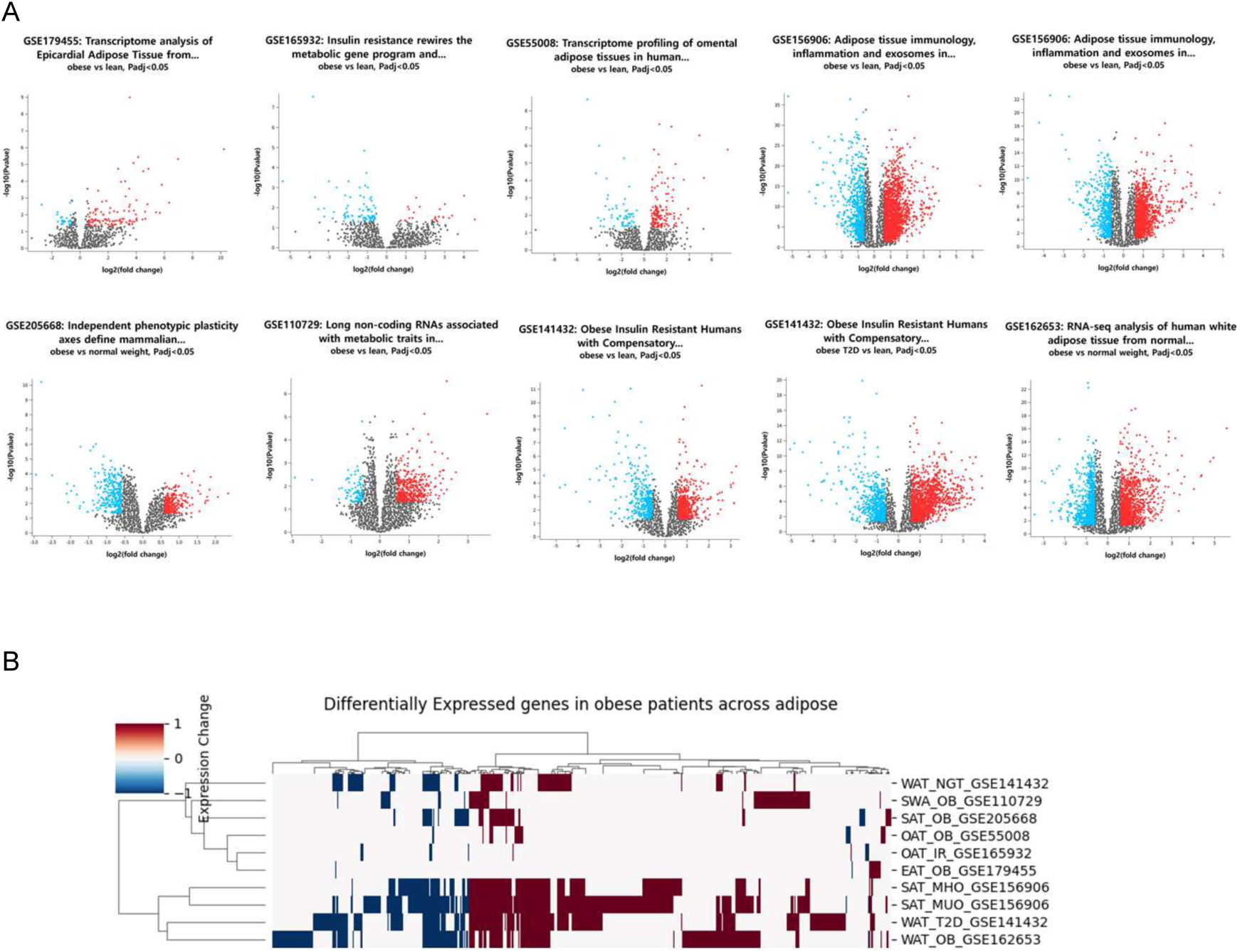
Differential Expression in Adipose Tissues by Obesity Status. **A.** Gene expression differences between adipose tissues from obese individuals and those with normal weight or lean mass are shown. Genes were identified as significantly up-regulated (red) or down-regulated (blue) based on an adjusted p-value threshold of <0.05 and a fold change cutoff of 1.5. **B.** A.Clustering of differentially expressed genes between studies.

**Figure S4.**
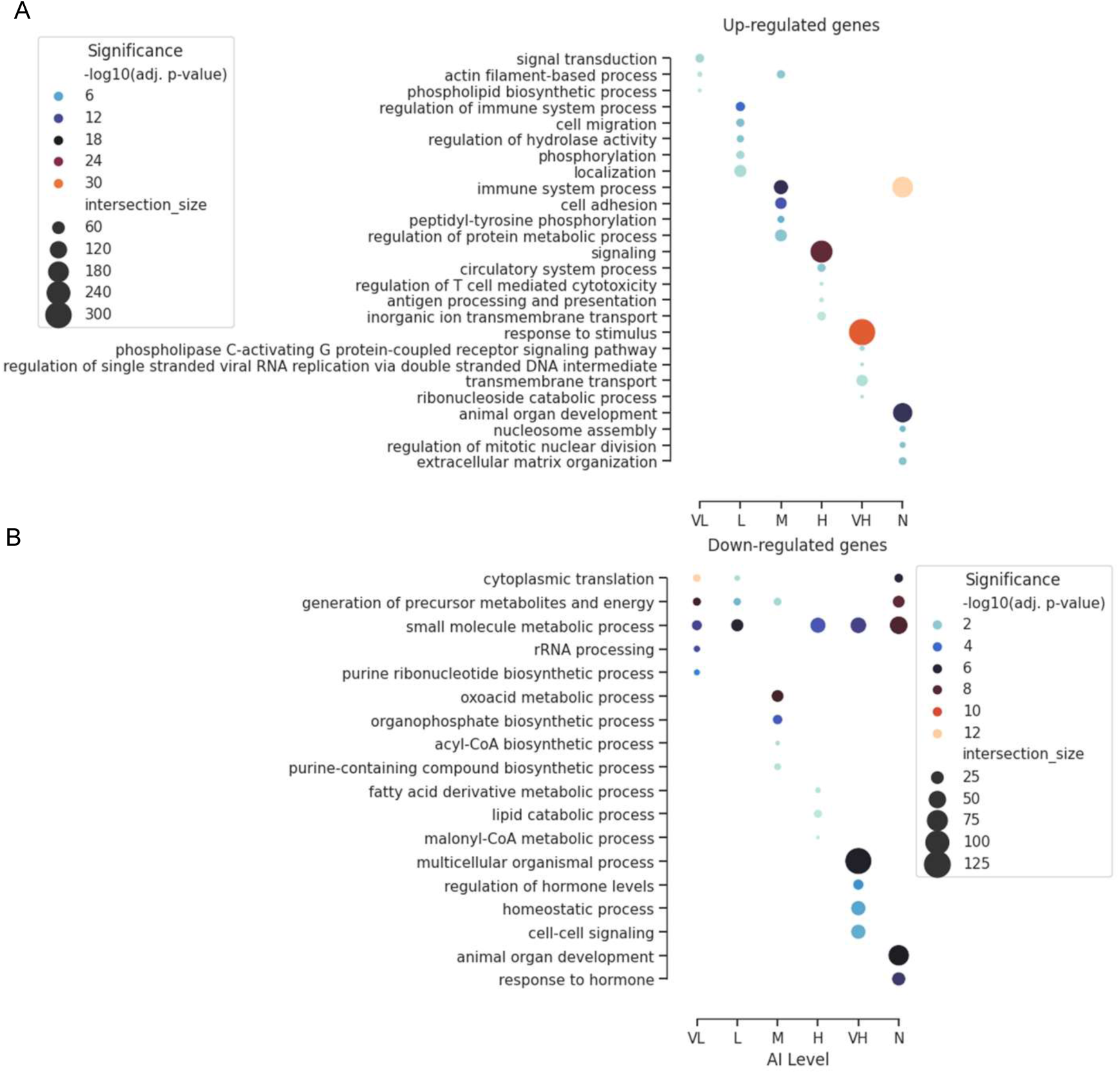
Biological processes enriched in differentially expressed eGenes related to obese adipose tissues. **A.** biological processes significantly enriched in each subset of up-regulated genes by allelic imbalance levels. **B.** biological processes significantly enriched in each subset of down-regulated genes by allelic imbalance level.

**Figure S5.**
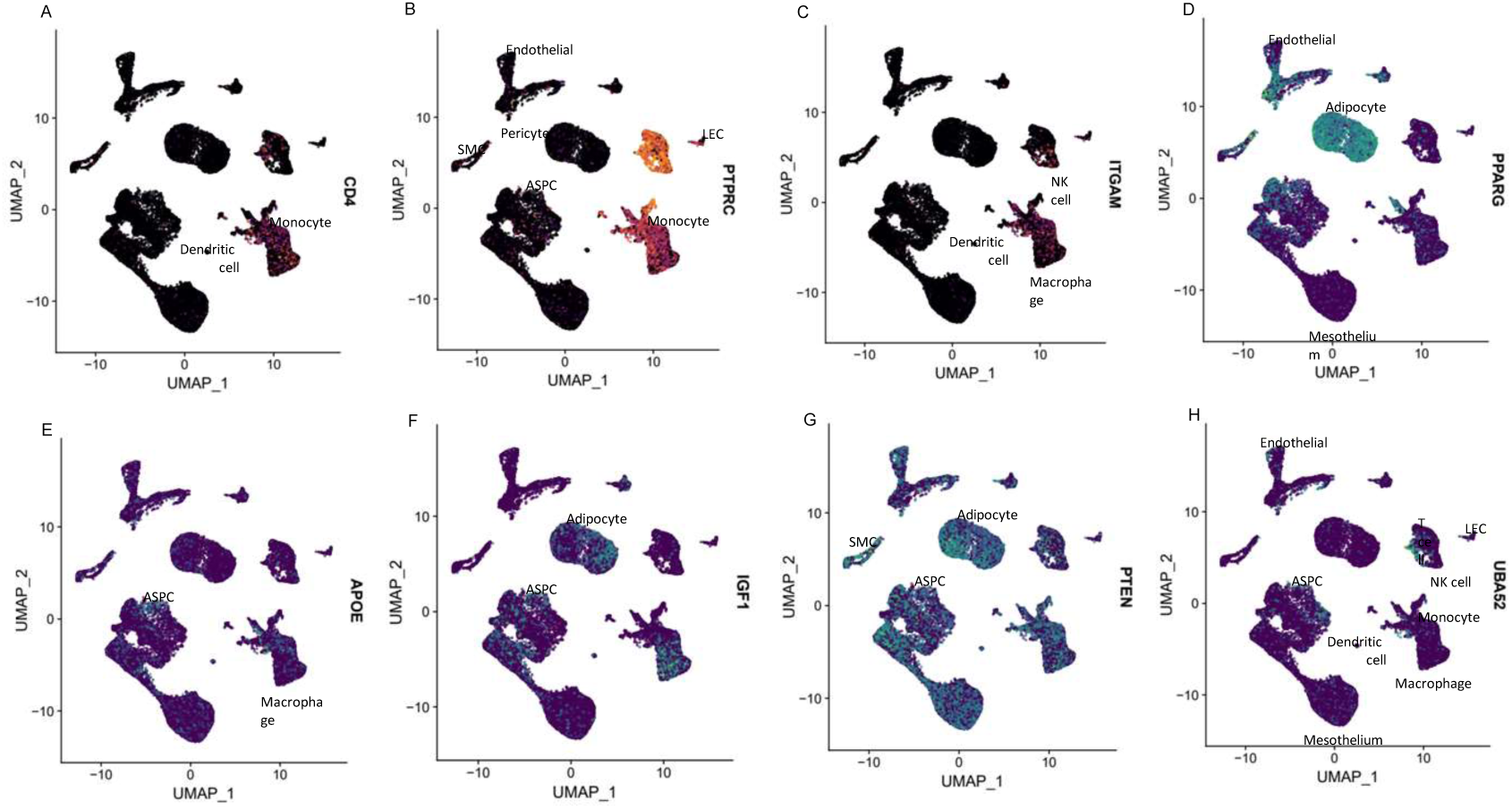
Cell specificity of differentially expressed genes in single cell transcriptome from human white adipose tissues. Gene expression at single cell level in human white adipose tissues for differentially expressed hub eGenes, including CD4, PTPRC, ITGAM, PPARG, APOE, IGF1, PTEN, and UBA52.

**Figure S6.**
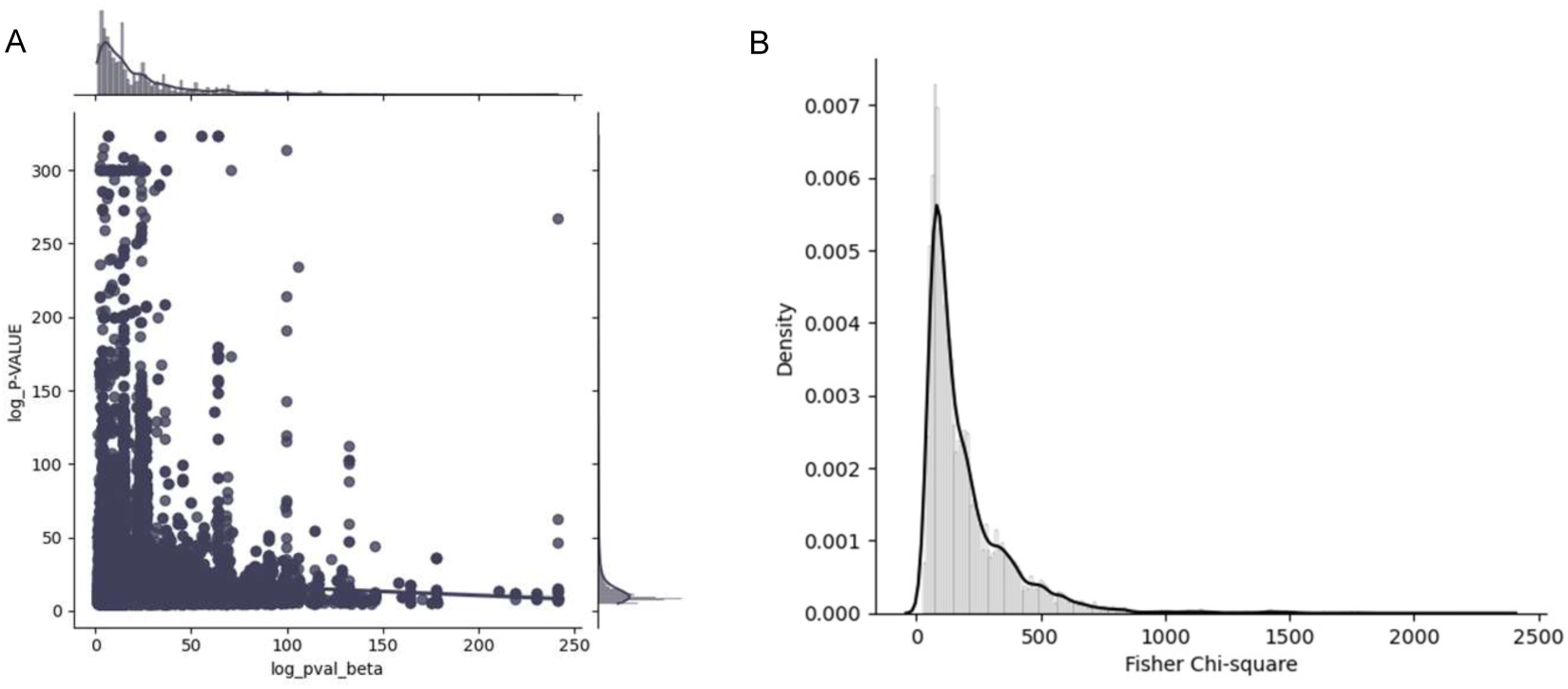
Composite probability for pairs of cis-eQTL and GWAS-SNP. **A.** Joint density distribution of p-values of cis-eQTL and GWAS-SNP that colocalized within DHS. **B.** Composite probability for those pairs of p-values using Fisher’s Method.

